# Seasonal dynamics of methane cycling microbial communities in Amazonian floodplain sediments

**DOI:** 10.1101/2020.05.04.076356

**Authors:** Júlia B. Gontijo, Andressa M. Venturini, Caio A. Yoshiura, Clovis D. Borges, José Mauro S. Moura, Brendan J. M. Bohannan, Klaus Nüsslein, Jorge L. Mazza Rodrigues, Fabiana S. Paula, Siu M. Tsai

## Abstract

The Amazonian floodplain forests are dynamic ecosystems of great importance for the regional hydrological and biogeochemical cycles and provide a significant contribution to the global carbon balance. Unique geochemical factors may drive the microbial community composition and, consequently, affect CH_4_ emissions across floodplain areas. Here we provide the first report of the *in situ* seasonal dynamics of CH_4_ cycling microbial communities in Amazonian floodplains. We asked how abiotic factors may affect both overall and CH_4_ cycling microbial communities and further investigated their responses to seasonal changes. We collected sediment samples during wet and dry seasons from three different types of floodplain forests, along with upland forest soil samples, from the Eastern Amazon, Brazil. We used high-resolution sequencing of archaeal and bacterial 16S rRNA genes combined with real-time PCR to quantify Archaea and Bacteria, as well as key functional genes indicative of the methanogenic (methyl coenzyme-M reductase – *mcr*A) and methanotrophic (particulate methane monooxygenase – *pmo*A) metabolisms. Methanogens were found to be present in high abundance in floodplain sediments and they seem to resist to dramatic seasonal environmental changes. Methanotrophs known to use different pathways to oxidise CH_4_ were detected, including anaerobic archaeal and bacterial taxa, indicating that a wide metabolic diversity may be harboured in this highly variable environment. The floodplain environmental variability, which is affected by the river origin, drives not only the sediment chemistry, but also the composition of the microbial communities. The results presented may contribute to the understanding of the current state of CH_4_ cycling in this region.

## Introduction

In the Amazon region, floodplain forests occupy an area of about 800,000 Km^2^ (Hess et al., 2015). These ecosystems comprise diversified and dynamic landscapes, which are exposed to seasonal flooding events by the expanding rivers, as a consequence of the periodic excessive rainfalls. Floodplains seem to play a significant role in the regional and global C budget (Junk, 1997; Moreira-Turcq, Seyler, Guyot & Etcheber, 2003; Pangala et al., 2017; Gedney, Huntingford, Comyn-Platt & Wiltshire, 2019). While uplands forests are considered important tropical methane (CH_4_) sinks, floodplains represent the largest natural sources of CH_4_ into the atmosphere (Conrad, 2009; Meyer et al. 2017; Gedney et al., 2019), including the significant process of CH_4_ transfer through trees (Pangala et al., 2017). Modelling studies have predicted that Amazonian floodplains may contribute up to 7% of total global CH_4_ emissions (Potter, Melack & Engle, 2014; Wilson et al., 2016).

In anoxic environments, the CH_4_ is generated as the final product of the anaerobic respiration by methanogenic archaea, which can use acetate, H_2_/CO_2_, formate, CO, or methylated compounds as substrates (Bridgham, Cadillo-Quiroz, Keller & Zhuang, 2013). The ability to use one or more substrates varies across the different methanogenic archaea taxa. Hence, substrate availability, along with other factors (biotic and abiotic), affects the diversity of these organisms in the environment (Barros et al., 2019).

In ecosystems that are sources of CH_4_, methanotrophs are particularly important for oxidising the CH_4_ and, therefore, attenuate net fluxes of this greenhouse gas into the atmosphere (Conrad, 2009; Ho et al., 2013). Aerobic methanotrophic bacteria can occur in terrestrial and aquatic environments, mainly at oxic/anoxic interfaces, where oxygen (O_2_) is available as an electron acceptor and CH_4_ as an energy and carbon source (Knief, 2015). In addition, the anaerobic oxidation of CH_4_ plays an important role in mitigating emissions of this greenhouse gas, and several types of this metabolism have been discovered in the recent years. However, the biochemical mechanisms performed by representatives of the Archaea (Nazaries, Murrell, Millard, Baggs, & Singh, 2013; Serrano-Silva, Sarria-Guzmán, Dendooven, & Luna-Guido, 2014) and Bacteria (Shen, Wu, Gao, Liu, & Li, 2016) are still vastly unknown (Scheller et al., 2020).

Despite the important role of the Amazonian floodplain forests for the global CH_4_ emissions (Sawakuchi et al., 2014, Gedney et al., 2019), the composition of their respective microbial communities remains mostly unexplored. A few studies have attempted to describe these communities (Conrad, Klose, Claus & Enrich-Prast, 2010; Conrad et al., 2011; Ji et al., 2016; Sawakuchi et al., 2016). Recently, using a microcosm experiment, Hernández et al. (2019) showed that methanogens are present in high abundance in Amazonian floodplain sediments regardless of the incubation conditions, while their abundance in upland forest soils increases only upon anaerobic incubation. Nevertheless, to date, there was no effort to characterise the seasonal dynamics of the methanogenic and methanotrophic groups *in situ*. In addition, the environmental variability present in floodplain ecosystems is likely to result in major differences in the CH_4_ cycling communities not only in temporal, but also in spatial scales. For instance, the riverine origin is a very important factor, as some waters may carry large amounts of inorganic suspensoids, such as the Amazon river, while others are comparatively poor in dissolved solids, such as the Tapajós river (Junk et al., 2011).

Here we provide the first in-depth characterisation of microbial communities responsible for CH_4_ cycling in Amazonian floodplain sediments and their seasonal dynamics *in situ*. We selected three floodplain areas with contrasting characteristics to explore part of the vast environmental variability found in this ecosystem. We asked how the abiotic factors inherent of each area may affect microbial community composition and further investigated season-driven changes. In addition, to examine the same seasonal dynamics in an area without flooding influences, an upland forest site was also studied. To tackle these questions, we used high-resolution sequencing of the 16S rRNA genes to assess archaeal and bacterial diversities, along with the quantification of key genes in the CH_4_ production (methyl coenzyme-M reductase – *mcr*A) and oxidation (particulate methane monooxygenase – *pmo*A). The results highlight the importance of the local environmental factors to drive both chemistry and microbiology of the floodplain forests. This information is of great relevance for the understanding of the current state of the regional CH_4_ cycle and to predict future scenarios under climate change influence.

## Methods

### Sediment and soil sampling

The studied sites are located in the region of Santarém and Belterra, in the central-western parts of the state of Pará, Brazil. The regional climate is classified as Am (Köppen), tropical humid, with a mean annual temperature of 26 ± 2 °C, and annual precipitation above 2,500 mm (Alvares, Stape, Sentelhas, Gonçalves & Sparovek, 2013). There are two well defined seasons, dry (DS; July – November) and wet (WS; December – June), with more than 70% of the rain concentrated in the latter.

Triplicate sediment samples were collected from three floodplain areas in May (WS) and October (DS) 2016, when the rivers reached maximum and minimum levels, respectively. These areas differed regarding their vegetation (Moura et al. 2008) and the adjacent river: FP1 (Floodplain 1 at Igarapé Jamaraquá, S2 49.077 W55 02.077), in the Tapajós river; FP2 (Floodplain 2 at Igarapé Maicá, S2 28.186 W54 38.831), in the Amazonas river; and FP3 (Floodplain 3 at Igarapé Açu, S2 22.747 W54 44.352), in the intersection of both rivers. In addition, we collected soil samples from an upland primary forest area, PFO (S2 51.326 W54 57.501), located in the Tapajós National Forest.

Sediment and soil samples were collected using a corer (5 cm diameter X 10 cm depth) and transported on ice to the laboratory. All samples were homogenised thoroughly and stored (−20 °C for DNA analyses and 4 °C for chemical analyses) and processed within two weeks. During the wet season, the water column in the floodplains ranged from 0.5 to 3 m. Dissolved oxygen (DO) and pH in the sediment-water interface were assessed using a YSI Professional Plus Instrument (Pro Plus, Yellow Springs, OH, USA). No water column was observed during the dry season sampling, however, site FP1 was water-logged.

### Chemical analyses

Sediment and soil samples were processed in the Laboratory of Chemical Analysis of the Department of Soil Science of the Luiz de Queiroz College of Agriculture (ESALQ/USP, Brazil), following procedures described by Camargo, Moniz, Jorge, & Valadares (2009). The following parameters were determined: pH in CaCl_2_; total nitrogen (N) by Kjeldahl method; phosphorus (P), potassium (K), calcium (Ca) and magnesium (Mg) by ion exchange resin extraction; sulphur (S) by calcium phosphate 0.01 mol L^-1^ extraction and turbidimetry determination; aluminium (Al) extraction by potassium chloride extraction 1 mol L^-1^; organic matter (OM) by the dichromate/titrimetric method; boron (B) by extraction with hot water; and the micronutrients copper (Cu), iron (Fe), manganese (Mn) and zinc (Zn) with a chelating agent, according to Lindsay and Norvell (1978).

### DNA extraction

The extraction of DNA from 0.25 g of sediment and soil samples was carried out in duplicate reactions using the PowerLyzer PowerSoil DNA Isolation Kit (MoBIO Laboratories Inc., Carlsbad, CA, USA), with an optimised protocol for tropical soils described by Venturini et al. (2020). Briefly, the adaptations included an extension in the vortex time to 15 min, followed by a 3 min centrifugation at 10,000 x g and incubation with C2 and C3 solutions at - 20 °C. DNA quantity and quality were assessed in 1% agarose gel and using a Nanodrop 2000c spectrophotometer (Thermo Fisher Scientific Inc., Wilmington, DE, USA) set for determining absorbance at the following wavelengths: 230, 260, 280 and 320 nm. Purified DNA samples were stored at -20 °C until processed.

### Archaeal and bacterial 16S rRNA gene sequencing

The diversity of archaeal and bacterial communities was assessed by high-throughput sequencing of the V4 region of 16S rRNA gene, using the following primer sets, respectively: 519f/915r (Klindworth et al., 2013; Stahl & Amman, 1991) and 515f/806r (Caporaso et al., 2011). Paired-end sequencing, with 2×250 bp reads, was performed in Illumina Hiseq 2500 platform, at Novogene Bioinformatics Technology (Beijing, China), using standard procedures.

### Quantitative PCR

Quantitative PCR (qPCR) was used to assess the abundance of archaeal and bacterial 16S rRNA genes, as well as functional gene markers for the methanogenic (methyl coenzyme-M reductase – *mcr*A) and methanotrophic (particulate methane monooxygenase – *pmo*A) metabolisms. Cycle conditions and primers used are described in the **Supporting Information Table S1**. The 10 μl reactions contained 5 μl of SYBR Green ROX qPCR (Thermo Fisher Scientific Inc., Wilmington, DE, USA), 0.2 μl of bovine serum albumin (Thermo Fisher Scientific Inc., Wilmington, DE, USA, 20 mg ml^-1^), 1 μl of each primer (5 pmol), 1 μl of DNA template (10 ng) and 1.8 μl of ultrapure water. Reactions were performed in triplicate using a StepOne Plus instrument (Applied Biosystems, Foster City, CA, USA). Gene abundance was estimated using a standard curve constructed with 10^0^ to 10^10^ copies of the targeted gene fragments amplified from the strains presented in **Supporting Information Table S1**.

### Bioinformatics and statistical analyses

All bioinformatics and statistical analyses were performed on R studio 3.5.1 (Rstudio Team, 2018). Raw sequences were analysed by inferring the amplicon sequence variants (ASVs) using the Dada2 1.9.3 package (Callahan et al. 2016). We obtained approximately 1.3 M sequences of Archaea and 4.1 M sequences of Bacteria. Reads with phred score > 30 were truncated at the positions 220 and 180 for Archaea and 200 and 190 for Bacteria. Sequences were error-corrected, dereplicated, merged and chimera-filtered. After quality control, 700,396 archaeal sequences with an average length of 383 bp, and 3,054,201 bacterial sequences with an average length of 253 bp were obtained. Taxonomy was assigned using the SILVA database (release 132, 12.13.2017). The ASV counts classified in the same taxonomic groups were summarised and normalised to within-sample relative abundance.

Statistical analyses and graphical visualisation were carried out using vegan 2.5-1 (Oksanen et al., 2018), ARTool 0.10.5 (Kay & Wobbrock, 2018), dunn.test 1.3.5 (Dinno, 2017), Hmisc 4.1-1 (Harrel, 2018), corrplot 0.84 (Wei & Simko, 2017) and ggplot2 3.1.0 (Wickham & Chang, 2016) packages. The methanogenic and methanotrophic taxa were filtered and grouped at the family level using the databases PhyMet2 (http://phymet2.biotech.uni.wroc.pl/) and Methanotroph Commons (http://www.methanotroph.org/wiki/taxonomy/), respectively. Shapiro-Wilk normality test and Levene’s homogeneity test were performed in order to define the most appropriate statistical test to be used to detect significant differences among treatments. Non-Metric Multidimensional Scaling (NMDS) and Analysis of Similarities (ANOSIM) were used to assess the similarities among samples regarding chemical properties (Euclidean distance) and community composition (Bray-Curtis distance). Envfit analysis was also carried out aiming to find the environmental variables related to the microbial community structure. Kruskal-Wallis with post-hoc Dunn’s test was used to determine statistical differences among the chemical properties of the studied areas. Two-way ANOVA of aligned rank transformed data (aiming to perform a non-parametric test using a parametric method) was used to investigate the effect of season and location on the relative abundance of CH_4_ cycling related taxa and functional gene quantities in the floodplains. The correlations among chemical properties, the relative abundance of taxonomic groups and qPCR data were performed and assessed using Spearman’s correlation coefficient.

## Results

### Seasonal changes in chemical properties

Lower values (p < 0.05) of N, P, K, Ca, Mg, Al, Cu, Mn, and Zn were observed for FP1, when compared to FP2 and FP3. Forest (PFO) soils presented higher OM and N contents in relation to the floodplains (**Supporting Information Table S2**). Multivariate analysis of the chemical profiles indicated significant differences among sites (**Figure 1**; **Supporting Information Table S3**). By contrast, no significant season effect was observed for any site.

**Figure 1.**
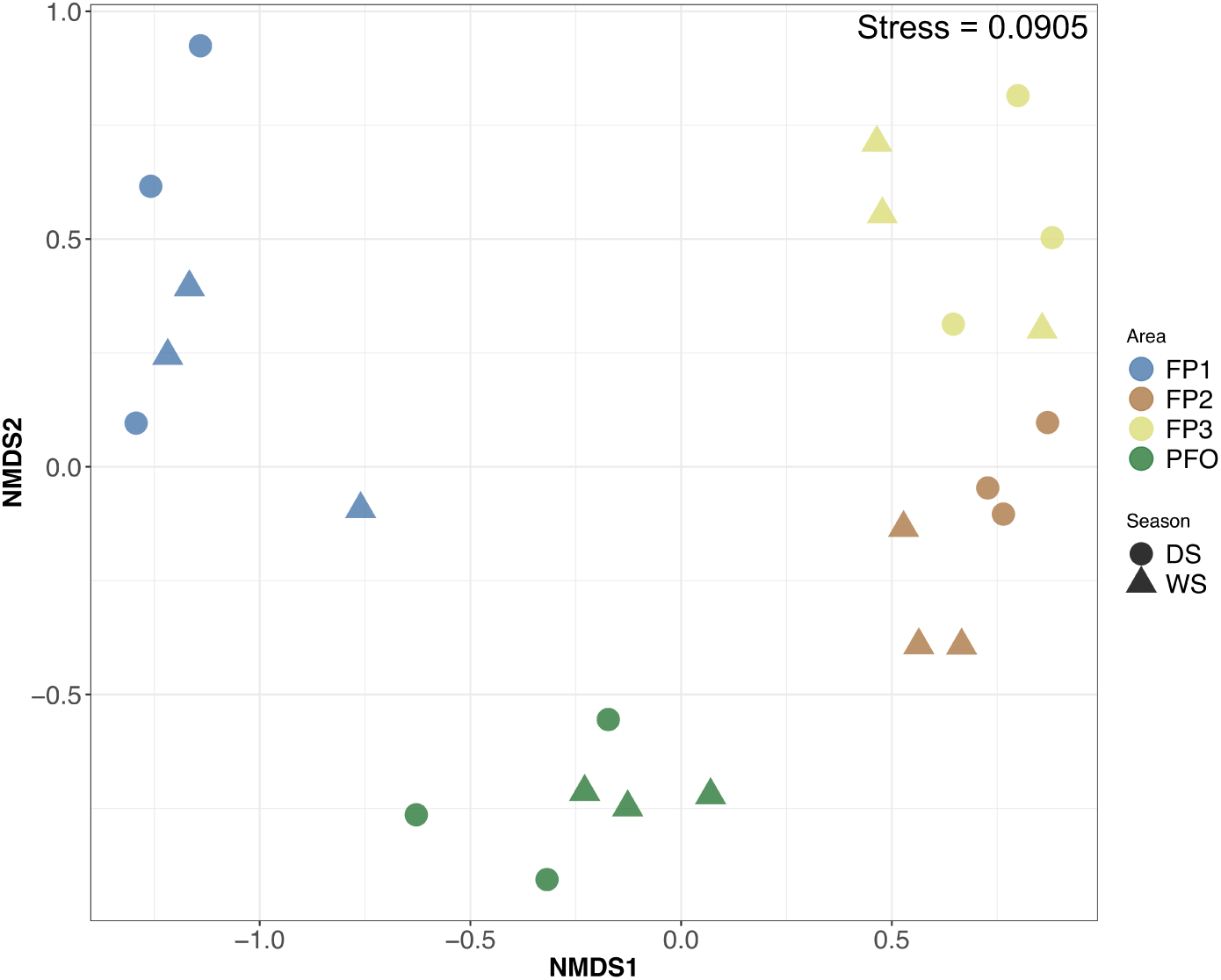
Clustering of the chemical properties of the floodplain sediments (FP1, FP2 and FP3) and upland forest soils (PFO) during wet (WS) and dry (DS) seasons. Plot is based on non-metric multidimensional scaling (NMDS) using the Euclidean distance index.

### Archaeal and bacterial communities

Analysis of similarities (Table 1) and NMDS ordination (**Figure 2a**) indicated no significant difference in the composition of the archaeal community among the three floodplain sites, albeit FP1 tended to form a separated cluster. pH and Mn correlated positively with the community composition in floodplain sites, while B, N and OM were correlated with forest soil communities (**Supporting Information Table S4**). Crenarchaeota and Thaumarchaeota were the dominant phyla in all floodplain sites (relative abundance average above 30%). Nanoarchaeota also presented high relative abundance in FP1 (average 16%), while Euryarchaeota were among the dominant phyla in FP2 and FP3, comprising the average of 18 and 26% of the archaeal communities, respectively. By contrast, major differences were observed between archaeal communities from forest soils and floodplain sediments. In the forest soils, the phylum Thaumarchaeota represented > 92% of the archaeal communities (**Supporting Information Figure S1a)**

**Table 1.** Analysis of Similarity of the microbial communities in the floodplain sediments and upland forest soils.

| Area | Archaea |  | Bacteria |  |
| --- | --- | --- | --- | --- |
|  | R | p-value | R | p-value |
| *All samples* | **0.6554** | **0.001** | **0.8925** | **0.001** |
| *Season comparison* |  |  |  |  |
| FP1 | -0.1852 | 0.900 | 0.8889 | 0.100 |
| FP2 | 0.2593 | 0.100 | 0.5926 | 0.100 |
| FP3 | 0.0000 | 0.900 | 0.1111 | 0.400 |
| PFO | -0.1852 | 0.900 | 0.1852 | 0.400 |
| *Area comparison* |  |  |  |  |
| FP1 vs. FP2 | 0.3148 | 0.057 | **0.8457** | **0.001** |
| FP1 vs. FP3 | 0.2809 | 0.055 | **0.8457** | **0.001** |
| FP2 vs. FP3 | 0.1698 | 0.140 | **0.6296** | **0.001** |
| FP1 vs. PFO | **0.6358** | **0.008** | **0.8025** | **0.001** |
| FP2 vs. PFO | **0.6790** | **0.002** | **0.7963** | **0.001** |
| FP3 vs. PFO | **0.6512** | **0.005** | **0.7037** | **0.004** |

FP1: Floodplain 1; FP2: Floodplain 2; FP3: Floodplain 3; PFO: Upland Forest. Bold values indicate statistical significance at p-value < 0.05. Distance index: Bray-Curtis.

**Figure 2.**
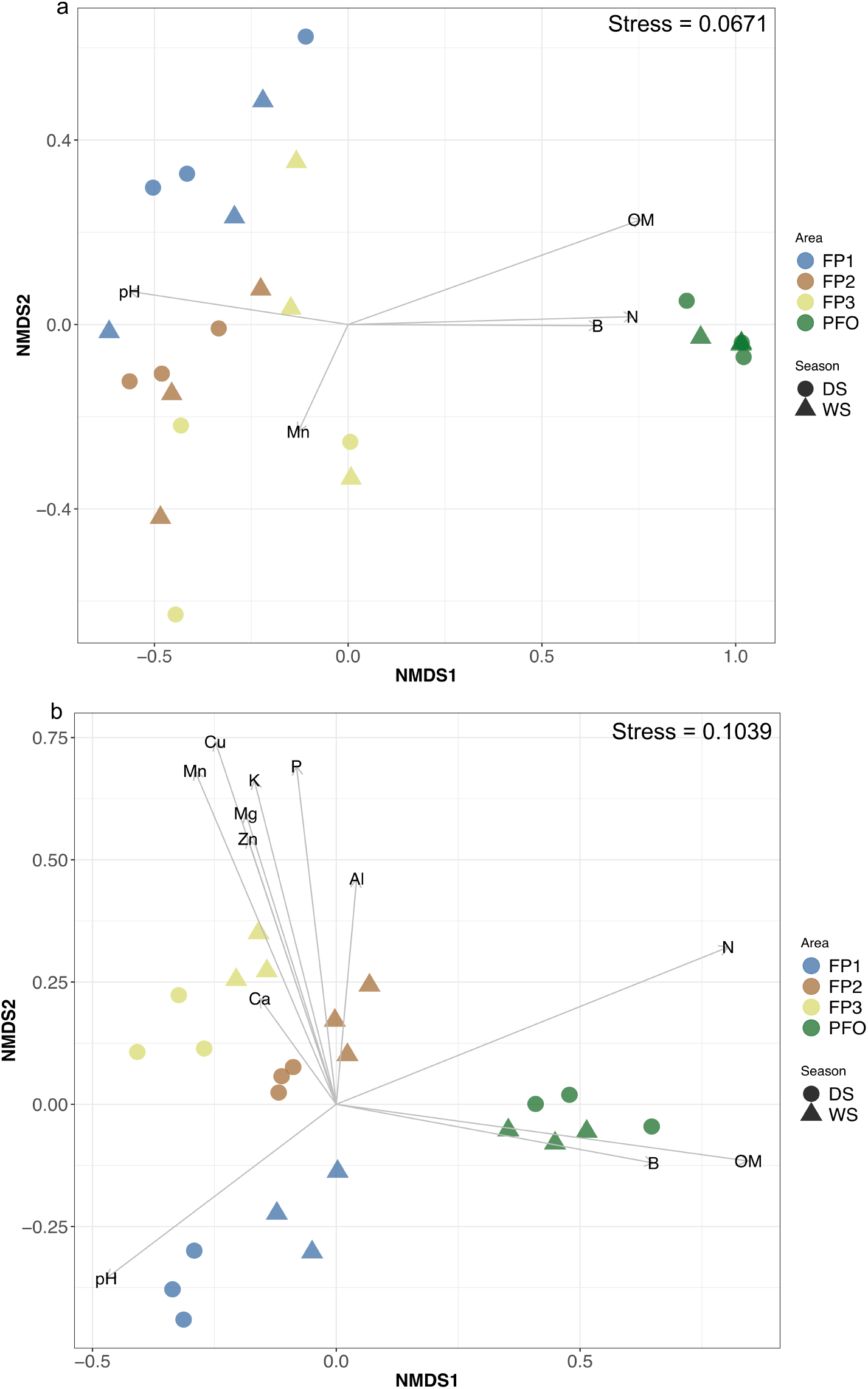
Clustering of the archaeal (a) and bacterial (b) communities in the floodplain sediments (FP1, FP2 and FP3) and upland forest soils (PFO) during wet (WS) and dry (DS) seasons, and correlation with environmental factors. Only environmental factors that share significant correlation (p < 0.05) with community structure are displayed with vectors: Al, aluminium; B: boron; Ca, calcium; Cu: cupper; K: potassium; Mg: magnesium; Mn: manganese; N: nitrogen; OM: organic matter; P: phosphorus; pH: hydrogen potential; Zn: zinc. Plot is based on non-metric multidimensional scaling (NMDS) using the Bray-Curtis distance index.

The composition of the bacterial communities was found to be significantly different across all sites **(Figure 2b, Table 1**). Although the bacterial communities from FP1 showed correlation only with pH, FP2 and FP3 were associated with increasing concentrations of Al, Ca, Mn, Zn, Mg, K, Cu and P (**Supporting Information Table S4**). Despite oscillations in relative abundance, the same phyla composed the dominant communities across all sites: Proteobacteria, Actinobacteria, Acidobacteria, Chloroflexi and Firmicutes (**Supporting Information Figure S1b**).

The abundance of archaea and bacteria in the samples was assessed by qPCR. Standard curves presented R^2^ values above 0.99 and amplification efficiencies between 75% and 99%. The average abundance (gene copies ng DNA^-1^) ranged from 3 × 10^3^ to 4 × 10^4^ for archaeal 16S rRNA, and 3 × 10^5^ to 1 × 10^6^ for bacterial 16S rRNA (**Supporting Information Figure S2**).

No seasonal effect on the overall microbial composition was observed for any site. By contrast, season was found to be a significant factor affecting bacterial abundance, while archaea responded to site x season interaction (**Table 2**).

**Table 2.** Two-way ANOVA of aligned rank transformed qPCR and sequencing data from the floodplains.

| Data | Area |  |  | Season |  |  | Area x Season |  |  |
| --- | --- | --- | --- | --- | --- | --- | --- | --- | --- |
|  | df | F | p | df | F | p | df | F | p |
| *Gene quantification* |  |  |  |  |  |  |  |  |  |
| 16S rRNA Archaea | 2, 12 | 3.745 | 0.055 | 1, 12 | 4.708 | 0.051 | 2, 12 | 4.401 | **0.037** |
| 16S rRNA Bacteria | 2, 12 | 3.843 | 0.051 | 1, 12 | 13.286 | **0.003** | 2, 12 | 0.406 | 0.675 |
| *mcrA* | 2, 12 | 1.388 | 0.287 | 1, 12 | 0.001 | 0.971 | 2, 12 | 0.180 | 0.837 |
| *pmoA* | 2, 12 | 0.013 | 0.987 | 1, 12 | 8.768 | **0.012** | 2, 12 | 0.056 | 0.945 |
| Ratio *mcrA:pmoA* | 2, 12 | 2.439 | 0.129 | 1, 12 | 3.314 | 0.094 | 2, 12 | 0.502 | 0.618 |
| *Methanogenic taxa* |  |  |  |  |  |  |  |  |  |
| Total | 2, 12 | 12.590 | **0.001** | 1, 12 | 0.676 | 0.427 | 2, 12 | 2.957 | 0.090 |
| Methanobacteriaceae | 2, 12 | 12.195 | **0.001** | 1, 12 | 0.439 | 0.520 | 2, 12 | 2.152 | 0.159 |
| Methanomassiliococcaceae | 2, 12 | 3.083 | 0.083 | 1, 12 | 1.961 | 0.187 | 2, 12 | 1.104 | 0.363 |
| Methanoperedenaceae | 2, 12 | 7.861 | **0.007** | 1, 12 | 12.009 | **0.005** | 2, 12 | 6.460 | **0.012** |
| Methanosarcinaceae | 2, 12 | 2.715 | 0.106 | 1, 12 | 10.269 | **0.008** | 2, 12 | 17.570 | **<0.001** |
| *Methanotrophic taxa* |  |  |  |  |  |  |  |  |  |
| Total | 2, 12 | 1.093 | **0.366** | 1, 12 | 0.240 | 0.633 | 2, 12 | 0.138 | 0.872 |
| Beijerinckiaceae | 2, 12 | 0.089 | 0.915 | 1, 12 | 0.125 | 0.729 | 2, 12 | 0.033 | 0.968 |
| Methylococcaceae | 2, 12 | 0.969 | 0.407 | 1, 12 | 2.618 | 0.132 | 2, 12 | 0.973 | 0.406 |
| Methylomonaceae | 2, 12 | 4.096 | **0.044** | 1, 12 | 8.133 | **0.015** | 2, 12 | 6.706 | **0.011** |
| NC10 | 2, 12 | 1.705 | 0.223 | 1, 12 | 0.245 | 0.630 | 2, 12 | 2.748 | 0.104 |

df: degrees of freedom, F: F-values, p: p-values. Bold values indicate statistical significance at p-value < 0.05.

### Methanogenic and methanotrophic communities

We searched the 16S rRNA gene sequence data for taxa with a reported role in CH_4_ production or consumption. The overall relative abundance of methanogenic archaea in the floodplain sediments varied widely, ranging from 2 to 34% on average, while it was below 1% in forest soils (**Figure 3a**). Methanobacteriaceae was the dominant methanogenic family across all samples, with relative abundance reaching 52% (31% on average). The relative abundance of Methanobacteriaceae and Methanoperedenaceae changed significantly with site location, but only Methanoperedenaceae and Methanosarcinaceae were significantly affected by season and by site x season interaction (**Table 2**). In addition, the class Bathyarchaeia, which has been suggested as a methanogenic taxon outside the Euryarchaeota phylum (Evans et al. 2015), was the main Crenarchaeota taxon detected in the floodplains, with a relative abundance ranging from 31 to 52% from the total archaeal community.

**Figure 3.**
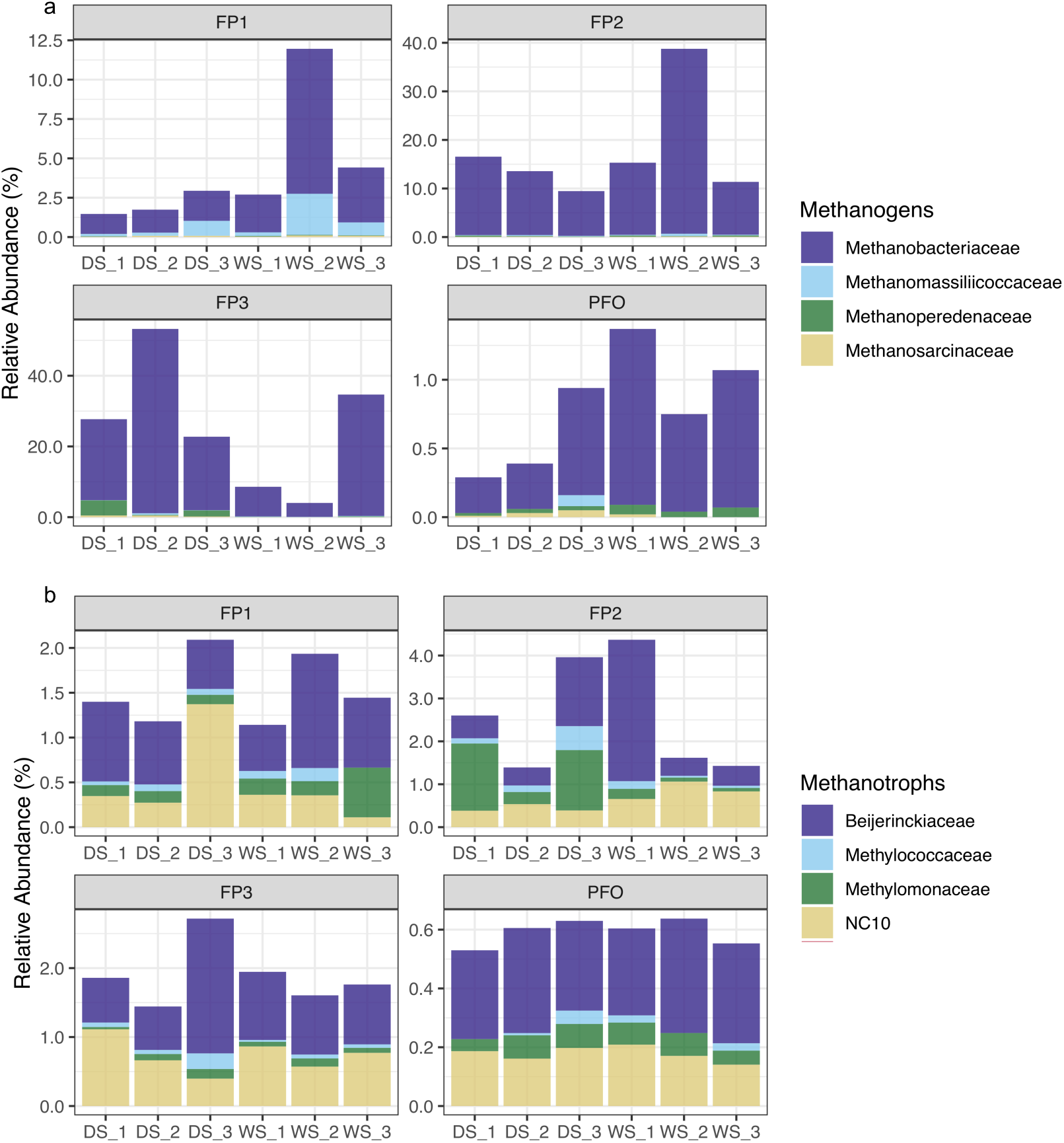
Relative abundance from of methanogenic (a) and methanotrophic (b) taxa in the floodplain sediments (FP1, FP2 and FP3) and upland forest soils (PFO) during wet (WS) and dry (DS) seasons.

Methanotrophic taxa represented only a small fraction of the bacterial community. The relative abundance of methanotrophs ranged from 1.5 to 2.7% in the floodplain sediments, while it was below 0.6% in the forest soils (**Figure 3b**). Groups from the Beijerinckiaceae family (Alphaproteobacteria class) and NC10 class (Rokubacteria phylum) were the most abundant methanotrophs in all areas, followed by Methylomonaceae and Methylococcaceae (Gammaproteobacteria class). Only Methylomonaceae changed significantly across sites and seasons, while all other methanotrophic families were not affected by both factors (**Table 2**).

When we assessed the abundance of the gene markers for methanogenesis (*mcr*A) and methanotrophy (*pmo*A), the patterns observed above were confirmed: in floodplain sediments, regardless of the season, there was a higher abundance of methanogens and methanotrophs in comparison to forest soil **(Figure 4a and 4b**). The *mcr*A gene abundance ranged from 2 × 10^3^ to 5 × 10^3^ copies ng DNA^-1^ in the floodplain sediments, while in the forest soils, it was below the detection limit. Gene counts for *pmo*A in floodplain sediments ranged from 1 × 10^2^ to 4 × 10^2^ copies ng DNA^-1^, while forest soils presented less than 5 × 10^1^ copies ng DNA^-1^. Considering only the floodplains, the abundance of the *mcr*A gene was not significantly affected by area or season, while *pmo*A varied with season (**Table 2**). This pattern was also observed in the *mcrA*:*pmoA* ratio, which tended to decrease in the dry season (**Table 2; Figure 4c)**.

**Figure 4.**
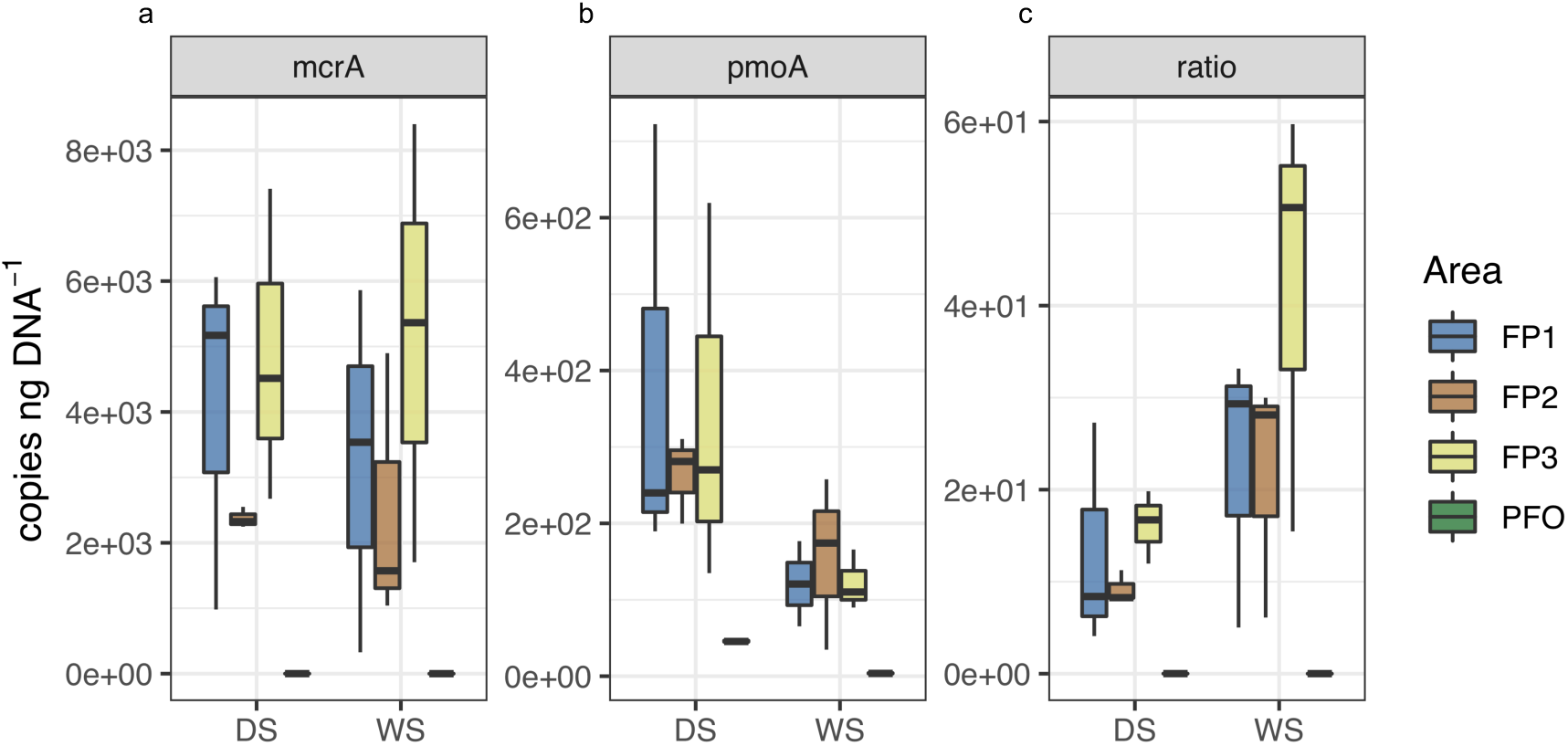
Number of copies per ng of DNA (copies ng DNA^-1^) of *mcr*A (a), *pmo*A (b) genes, and *mcr*A:*pmo*A ratio (c) in the floodplain sediments (FP1, FP2 and FP3) and upland forest soils (PFO) during wet (WS) and dry (DS) seasons.

To search for correlations between CH_4_ cycling microbes and abiotic factors in the floodplains, we used the Spearman coefficient. **Figure 5** shows significant (p < 0.05) positive and negative correlations observed for floodplain areas across seasons. The *pmo*A gene abundance was strongly correlated with DO during the wet season. The pH value was positively correlated with the relative abundance of Methanomassiliicoccaceae, and negatively correlated with Methanobacteriaceae, Methanoperedenaceae and NC10. Methylomonaceae was positively correlated with OM, while Methanobacteriaceae and Methanoperedenaceae correlated with N and P. Interestingly, Methanoperedenaceae also showed a positive correlation with Mn contents.

**Figure 5.**
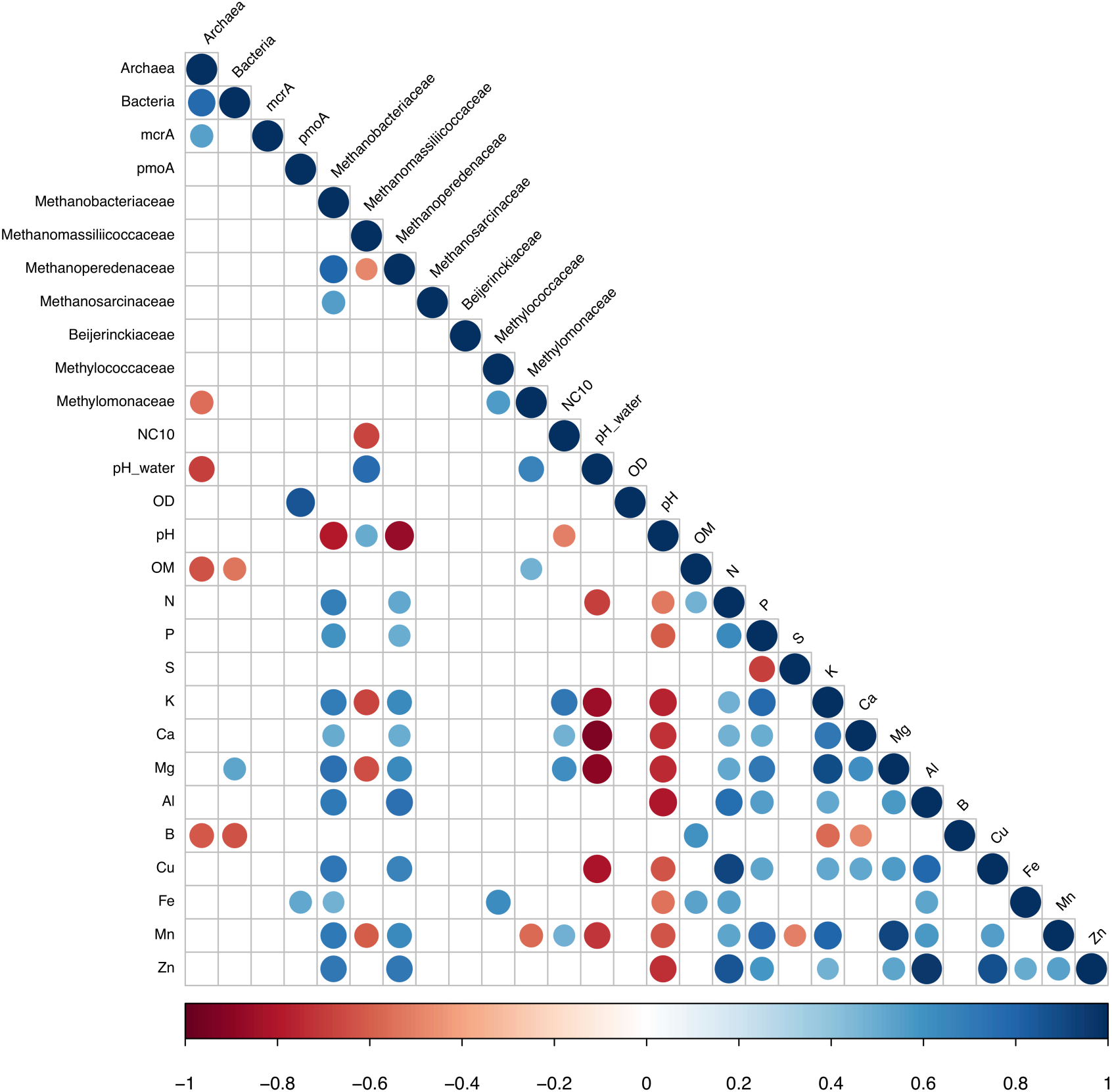
Spearman’s correlations between the abundance of 16S rRNA and CH_4_ marker genes, methanogenic and methanotrophic taxa and chemical properties in the floodplain (FP1, FP2 and FP3) sediments. Significant correlations (p<0.05) are marked in blue (positive) and in red (negative). Circle size is proportional to the R^2^ value.

## Discussion

The Amazon river, termed “white-water”, transports large amounts of nutrient-rich sediments, which are carried to the floodplain forests during the raining season. By contrast, the Tapajós river, termed “clear-water”, delivers low amounts of sediments and dissolved solids (Junk et al., 2011). The transfer of materials by the Amazon river to sediments at sites FP2 and FP3 was indicated by their higher contents of trace metals, such as Cu, Mn and Zn. As a consequence, these sites presented a higher chemical similarity to each other, when compared to FP1, which is exposed exclusively to the Tapajós river. In fact, FP1 clustered separately from the other floodplains, suggesting that due to the low nutrient content of the Tapajos, forest-related factors may have a stronger influence on the chemical composition of these sediments.

In the Amazon region, several studies already showed the importance of soil chemical properties on the soil microbial community composition (Rodrigues et al. 2013) and function (Paula et al. 2014; Lammel, Feigl, Cerri, & Nüsslein, 2015). Here we show that archaeal and bacterial communities in sediments from the three floodplains analysed are significantly distinct from upland forest soil communities regarding their composition. In addition, we observed major differences among floodplain areas with distinct environmental characteristics. This information is valuable to detangle which environmental factors drive the variations in microbe-driven ecosystem processes. It is interesting that the archaeal and bacterial communities from the FP1 did not cluster together with upland forest soil communities. This might indicate that flooding is a major determinant of the composition of the microbial communities in the floodplain forests, and also that environmental factors not assessed in this study are likely to be affecting these communities.

Our data indicated that the composition of the microbial communities did not change with season, although this factor affected the abundance of archaea and bacteria. Microbes are known for their high degree of metabolic flexibility and physiological adaptations that may allow them to endure under changing environmental conditions (Meyer, Lipson, Martín, Schadt, Schmidt, 2004; Ye et al., 2018). The periodic flooding events may favour microbial taxa that are adapted to these environmental oscillations.

Upland forest soils were largely dominated by the phylum Thaumarchaeota. By contrast, in the floodplains, Euryarchaeota and Crenarchaeota were also among the dominant groups. To date, all isolates of methanogenic archaea belong to the phylum Euryarchaeota (Evans et al., 2019). However, recent metagenomic studies have suggested that the class Bathyarchaeia (representing > 99% of Crenarchaeota in our study) may have a potential function in CH_4_ production and consumption (Evans et al., 2015). The role of Bathyarchaeia in the CH_4_ cycle is yet to be clarified, but the presence of genes related to other steps in the anaerobic degradation of organic matter in their genomes, including acetogenesis (He et al. 2016), indicates that this group may have an important role in anaerobic environments, such as floodplain sediments.

We also observed a higher relative abundance of the class Wosearchaeia (representing > 99% of Nanoarchaeota) in FP1, in relation to the other floodplain areas. Although scant information is available for this class, studies have suggested that some groups may establish syntrophic interactions with methanogens and provide metabolic complementation (Liu et al., 2018). This might indicate a potential role of this group in the CH_4_ cycle of FP1, which has oligotrophic characteristics, if compared to the other floodplains. Together, Euryarchaeota, Bathyarchaeia and Woesearchaeia represent more than half of the total archaeal community in the floodplains during all seasons, highlighting the importance of the CH_4_ metabolism in the biogeochemistry of these environments.

Among the methanogens belonging to the Euryarchaeota, Methanobacteriaceae was the dominant family in all areas. This group is widely distributed in anaerobic environments across the globe and is known to produce CH_4_ mainly through the hydrogenotrophic pathway (Evans et al., 2019). Methanosarcinacea, which accounted only for a small fraction of the methanogens, is a metabolically versatile group and is able to utilise H_2_/CO_2_, acetate, methylamines, and methanol as substrates for methanogenesis (Welander & Metcalf, 2008; Fournier, 2009; Evans et al.; 2019). Methanomassiliicoccaceae, which was part of the dominant groups only in FP1, is known as an obligately methylotrophic and hydrogen-dependent methanogen (Nkamga & Drancourt, 2016). In our study, we observed a prevalence of microbes with the potential to perform the hydrogenotrophic pathway. However, investigations on the RNA level and on the isotopic signal would be required to determine the dominant metabolic route in the Amazonian floodplains. Transcriptional data will also be essential to explain the emission profiles reported for these ecosystems (Angle et al., 2017). At the DNA level, we did not find significant variations in both relative and absolute abundance of methanogens between both seasons. Despite the lack of knowledge in the mechanisms underpinning this process, the resistance of methanogens to drainage was also reported by Ma and Lu (2011). It is likely that those microbes may present protection against oxidative stress, as already reported for methanogens (Angle et al., 2017; Lyu, Shao Akinyemi, & Whitman, 2018). According to Hernández et al. (2019), the abundance of methanogens could be an indication of the flooding history of the Amazon floodplain. In a microcosm experiment, the authors observed the survival of the methanogenic populations to short, but not to long periods of desiccation.

Regarding the CH_4_-oxidising microbes, the Type II-b methanotrophs from the Beijerinckiaceae family were found to compose large fractions of the communities and did not seem to be affected by season. These organisms are known to endure under fluctuations in the environment, such as variable O_2_ and CH_4_ availability. Their stability is owed to several strategies, including dormancy (Eller, Krüger, & Frenzel, 2005; Krause, Lüke & Frenzel, 2012; Ho et al., 2013) and ability to use different carbon substrates (Dedysh, Knief, & Dunfield, 2005; Dedysh, Haup, & Dunfield, 2016). Such versatility could confer an advantage in the floodplain environment, which may present not only temporal, but also spatial variation in CH_4_ availability (Moura et al., 2008). Other groups detected at lower relative abundances, the Methylococcaceae and Methylomonaceae, can use the ribulose monophosphate pathway, common to all Type I-a methanotrophs (Knief, 2015). In general, members of the Type I of methanotrophic bacteria are very responsive to high substrate availability, but have their abundance reduced quickly under O_2_ limitation or other adverse conditions, as the contrasting wet and dry seasons in the Amazon basin. The abundance of methanotrophs (assessed by quantifying the *pmo*A gene), was affected by seasonality and positively correlated with dissolved O_2_ during the wet season. The *pmo*A gene is present in all known Type I metabolic types and also in the anaerobic denitrifying NC10, while it can be present also in some Type II taxa with exceptions among Beijerinckiaceae (Knief, 2015).

The rate of CH_4_ emission is related to the balance between the activity of methanogens and methanotrophs in the environment (Malyan et al., 2016). Although many studies showed that Amazonian floodplains are an important source of CH_4_ (Ringeval et al., 2014; Potter et al., 2014; Barbosa et al., 2020), Koschorreck (2000) demonstrated that the CH_4_ emission rates may decrease to zero when the sediments become exposed to air. In fact, we observed a decrease of *mcr*A:*pmo*A ratios during the dry season in all areas. The drainage of the floodplains allows for the oxygenation of the sediment, which could promote an increase in abundance and/or activity of the aerobic methanotrophs in relation to the methanogenic community.

Aerobic CH_4_-oxidising microorganisms have long been recognised to play an important role in the regulation of CH_4_ emissions to the atmosphere (Graf et al. 2018). However, in the recent years, anaerobic oxidation of CH_4_ (AOM) has been shown to be more widespread than previously predicted in areas of high organic matter availability (Evans et al., 2019). In the floodplain sediments, we detected both archaeal and bacterial taxa with the reported capability to carry out anaerobic CH_4_ oxidation. For instance, the methanogenic family Methanoperedenaceae (Timmers et al. 2017) has been proposed as a potential anaerobic methanotrophic archaeal (ANME) group, using the nitrate-dependent reverse methanogenesis (Yan & Ferry, 2018) or in consortia with sulphate-reducing bacteria (Su et al., 2019). Here we found a positive correlation between Metanoperedenaceae relative abundance and Mn contents of the sediment. This could suggest a potential role of these microbes, as it has been demonstrated that members of the ANME group, including the Methanoperedenaceae, can be coupled with the dissimilatory reduction of metals, including Mn (Scheller, Yu, Chadwick, McGlynn, & Orphan, 2016, Leu et al., 2020). The bacterial class NC10, which is among the dominant methanotroph groups detected in this study, has been reported to contain groups that are able to oxidise CH_4_ under anaerobic conditions using nitrite as an electron acceptor, a process that generates oxygen, which is subsequently used to oxidise CH_4_ as carbon source (Ettwig et al., 2010; Padilla et al., 2016). This capability provides an advantage for NC10 to thrive in oxygen-depleted environments (Shen et al., 2016). The metabolic diversity and the biogeography of AOM are still largely unknown (Cui, Ma, Qi, Zhuang, & Zhuang, 2015) and studies providing insights in this field are in high demand. Recently, Gabriel et al. (2020) incubated Amazonian floodplains sediments in anaerobic reactors and observed the presence of AOM coupled with iron reduction. Here we report the *in situ* presence of archaeal and bacterial taxa with the potential to oxidise CH_4_ anaerobically in Amazonian floodplains.

We presented the first report on the *in situ* seasonal dynamics of CH_4_ cycling microbial communities in three different types of floodplains located in the Eastern Amazon. We observed that methanogens are present in high abundance and they seem to resist the dramatic environmental changes that occur between seasons. Methanotrophs that use different pathways to oxidise CH_4_ were detected, indicating that a wide metabolic diversity may be harboured in this highly variable environment. This environmental variability, which is remarkably affected by the river origin, drives not only the floodplain sediment chemistry, but also the composition of the microbial communities. In the light of climate change, the data provided in this study may contribute to the understanding of the current state of the CH_4_ cycling in the Amazonian floodplains, which is essential to predict future scenarios.

## Data Accessibility

16S rRNA sequence data were deposited on NCBI’s Sequence Read Archive (SRA) under the accession number PRJNA629547.

## Supporting information

supporting information Gontijo et al

## Author Contributions

JBG, SMT, BJMB, KN and JLMR designed the research; JBG, AMV, JMSM, and CDB collected the samples; JBG and AMV performed the bench work; JBG and CAY analysed the data; JBG, FSP and SMT wrote the manuscript; all authors contributed to the final manuscript version.

## Acknowledgments

This study was financed in part by the São Paulo Research Foundation (FAPESP grants 2014/50320-4, 2015/13546-7, 2017/26138-0, 2018/14974-0 and 2019/25931-3), National Council for Scientific and Technological Development (CNPq grants 133769/2015-1 and 311008/2016-0), and the Coordination for the Improvement of Higher Education Personnel - Brasil (CAPES) - Finance Code 001. We especially thank Wagner Piccinini, Erika B. Cesar, Liana C. Rossi and the ECOFOR and LBA teams for their contribution to the field expeditions.

