## supporting information Gontijo et al for "Seasonal dynamics of methane cycling microbial communities in Amazonian floodplain sediments"

**Table S1.** Real-time PCR conditions used in this study.

| Genes | Standard DNA | Primers | Reference | Amplifications conditions |
|---|---|---|---|---|
| *Taxonomic genes* | | | | |
| 16S rRNA Archaea | DSM 23604 *Methanolinea mesophila* | ARC787f ARC1059r | Yu et al. (2005) | 95 ºC - 10 min; 45 cycles, 95 ºC - 45 s, 57 ºC - 45 s, 72 ºC - 45 s; 72 ºC - 10 min |
| 16S rRNA Bacteria | DSM 11192 *Gordonia* sp. | 926f 1062r | Gregoris et al. (2011) | 95 ºC - 15 min; 40 cycles, 95 ºC - 45 s, 59 ºC - 15 s, 72 ºC - 20 s; 72 ºC - 5 min |
| *$CH_4$ cycle marker genes* | | | | |
| *mcr*A | DSM 23604 *Methanolinea mesophila* | mlas mcrA-rev | Steinberg & Regan (2008) | 95 ºC - 4 min; 40 cycles, 95 ºC - 30 s, 60 ºC - 45 s, 72 ºC - 30 s; 72 ºC - 10 min |
| *pmo*A | DSM 17706 Methylosinus sporium | A189f MB661r | Holmes et al. (1995) Costello & Lidstrom (1999) | 95 ºC - 4 min; 40 cycles, 95 ºC - 30 s, 58 ºC - 45 s, 72 ºC - 45 s; 72 ºC - 10 min |

**Table S2.** Chemical properties of the floodplain sediments and upland forest soils during wet and dry seasons.

| Chemical Properties | Units | FP1 | | FP2 | | FP3 | | PFO | |
|---|---|---|---|---|---|---|---|---|---|
| | | WS | DS | WS | DS | WS | DS | WS | DS |
| pH water | - | 6.85 ± 0.04$^{a}$ | - - | 6.39 ± 0.19$^{b}$ | - - | 6.26 ± 0.14$^{b}$ | - - | - - | - - |
| DO water | mg $L^{-1}$ | 3.29 ± 0.60$^{a}$ | - - | 4.34 ± 0.90$^{a}$ | - - | 3.48 ± 0.12$^{a}$ | - - | - - | - - |
| pH | - | 4.27 ± 0.15$^{aA}$ | 4.40 ± 0.26$^{aA}$ | 3.97 ± 0.06$^{bA}$ | 3.80 ± 0.00$^{bB}$ | 4.07 ± 0.06$^{abA}$ | 3.70 ± 0.10$^{bB}$ | 3.57 ± 0.06$^{cA}$ | 3.20 ± 0.17$^{cB}$ |
| OM | g $dm^{-3}$ | 16.67 ± 7.77$^{bA}$ | 5.33 ± 2.31$^{cB}$ | 16.67 ± 7.64$^{bA}$ | 12.33 ± 0.58$^{bA}$ | 7.33 ± 1.15$^{bA}$ | 6.67 ± 2.08b$^{cA}$ | 44.33 ± 0.58$^{aA}$ | 39.33 ± 15.37$^{aA}$ |
| N | g $Kg^{-1}$ | 0.65 ± 0.31$^{cA}$ | 0.28 ± 0.08$^{cB}$ | 1.53 ± 0.18$^{bA}$ | 1.40 ± 0.07$^{bA}$ | 1.35 ± 0.27$^{bA}$ | 1.04 ± 0.05$^{bB}$ | 3.25 ± 0.41$^{aA}$ | 2.77 ± 0.81$^{aA}$ |
| P | mg $dm^{-3}$ | 3.67 ± 1.15$^{bA}$ | 2.67 ± 1.53$^{cA}$ | 12.00 ± 2.65$^{aA}$ | 14.67 ± 2.08$^{aA}$ | 14.67 ± 4.04$^{aA}$ | 14.00 ± 4.00$^{aA}$ | 6.67 ± 0.58$^{bB}$ | 9.00 ± 1.00$^{bA}$ |
| S | mg $dm^{-3}$ | 23.67 ± 25.15$^{aA}$ | 14.00 ± 8.19$^{aA}$ | 13.33 ± 16.17$^{aA}$ | 11.67 ± 0.58$^{aA}$ | 13.00 ± 12.29$^{aA}$ | 12.67 ± 6.03$^{aA}$ | 6.33 ± 1.53$^{aB}$ | 19.33 ± 4.04$^{aA}$ |
| K | mmolc $dm^{-3}$ | 0.13 ± 0.06$^{cA}$ | 0.13 ± 0.06$^{cA}$ | 0.87 ± 0.47$^{abB}$ | 1.77 ± 0.38$^{abA}$ | 1.97 ± 0.06$^{aA}$ | 2.07 ± 0.35$^{aA}$ | 0.57 ± 0.06$^{bB}$ | 0.87 ± 0.06$^{bA}$ |
| Ca | mmolc $dm^{-3}$ | 3.00 ± 0.00$^{cA}$ | 4.33 ± 2.31$^{bA}$ | 5.00 ± 0.00$^{aB}$ | 11.00 ± 2.00$^{aA}$ | 5.67 ± 1.53$^{aA}$ | 7.33 ± 2.52$^{abA}$ | 3.67 ± 0.58$^{bA}$ | 3.33 ± 0.58$^{bA}$ |
| Mg | mmolc $dm^{-3}$ | 1.00 ± 0.00$^{bA}$ | 1.00 ± 0.00$^{bA}$ | 3.00 ± 0.00$^{aA}$ | 3.33 ± 0.58$^{aA}$ | 5.33 ± 1.53$^{aA}$ | 7.00 ± 1.00$^{aA}$ | 3.00 ± 1.00$^{aA}$ | 1.67 ± 0.58$^{bB}$ |
| Al | mmolc $dm^{-3}$ | 5.33 ± 4.16$^{cA}$ | 2.67 ± 1.53$^{bA}$ | 35.33 ± 8.62$^{aA}$ | 27.00 ± 2.00$^{aA}$ | 16.00 ± 1.73$^{bA}$ | 27.67 ± 16.56$^{aA}$ | 20.67 ± 1.53$^{bA}$ | 15.67 ± 2.52$^{aA}$ |
| B | mg $dm^{-3}$ | 0.33 ± 0.06$^{bA}$ | 0.20 ± 0.01$^{bB}$ | 0.40 ± 0.10$^{abA}$ | 0.23 ± 0.06$^{bB}$ | 0.20 ± 0.00$^{bA}$ | 0.19 ± 0.03$^{bA}$ | 0.70 ± 0.00$^{aA}$ | 0.45 ± 0.11$^{aB}$ |
| Cu | mg $dm^{-3}$ | 0.23 ± 0.06$^{bA}$ | 0.10 ± 0.00$^{bB}$ | 3.53 ± 0.25$^{aA}$ | 3.10 ± 0.17$^{aB}$ | 3.53 ± 1.38$^{aA}$ | 2.80 ± 0.17$^{aA}$ | 0.40 ± 0.17$^{bA}$ | 0.27 ± 0.06$^{bA}$ |
| Fe | mg $dm^{-3}$ | 98.00 ± 59.57$^{aA}$ | 111.00 ± 67.45$^{bA}$ | 145.67 ± 24.34$^{aB}$ | 286.33 ± 54.52$^{aA}$ | 118.67 ± 9.29$^{aA}$ | 129.67 ± 5.69$^{bA}$ | 150.00 ± 6.93$^{aA}$ | 166.67 ± 59.48$^{abA}$ |
| Mn | mg $dm^{-3}$ | 0.50 ± 0.26$^{cA}$ | 0.30 ± 0.00$^{dA}$ | 64.80 ± 5.70$^{aA}$ | 26.67 ± 2.91$^{bB}$ | 115.10 ± 62.73$^{aA}$ | 107.23 ± 22.92$^{aA}$ | 4.00 ± 0.96$^{bA}$ | 2.07 ± 0.85$^{cB}$ |
| Zn | mg $dm^{-3}$ | 0.20 ± 0.10$^{bA}$ | 0.10 ± 0.00$^{bA}$ | 5.60 ± 0.75$^{aA}$ | 4.83 ± 0.70$^{aA}$ | 3.03 ± 1.19$^{aA}$ | 3.27 ± 1.76$^{aA}$ | 0.27 ± 0.06$^{bA}$ | 0.17 ± 0.06$^{bB}$ |

pH water: hydrogen potention of the water; DO water: dissolved oxygen of the water; pH: hydrogen potential; OM: organic matter; N: nitrogen, P: phosphorus; S: sulphur; K: potassium; Ca: calcium; Mg: magnesium; Al: aluminium; B: boron; Cu: cupper; Fe: iron; Mn manganese; Zn: zinc. Values are presented as means ± standard deviation. Lower-case letters indicate the comparison between areas in the same season and upper-case letters indicate the comparison between seasons for each area. Sites not sharing the same letter are significantly different from each other (Kruskal-Wallis and Dunn Test; $p < 0.05$). FP1: Floodplain 1; FP2: Floodplain 2; FP3: Floodplain 3; PFO: Upland Forest. WS: Wet season; DS: Dry season.

**Table S3.** Analysis of similarity of chemical profiles of floodplain sediments and forest soils.

| Area | Chemical Properties | |
|---|---|---|
| | R | p value |
| *All samples* | 0.8763 | **0.001** |
| *Season comparison* | | |
| FP1 | 0.1481 | 0.400 |
| FP2 | 0.5926 | 0.100 |
| FP3 | 0.1852 | 0.300 |
| PFO | 0.6667 | 0.100 |
| *Area comparison* | | |
| FP1 vs. FP2 | 0.7932 | **0.001** |
| FP1 vs. FP3 | 0.7130 | **0.003** |
| FP2 vs. FP3 | 0.6636 | **0.001** |
| FP1 vs. PFO | 0.7623 | **0.002** |
| FP2 vs. PFO | 0.8642 | **0.001** |
| FP3 vs. PFO | 0.7963 | **0.001** |

FP1: Floodplain 1; FP2: Floodplain 2; FP3: Floodplain 3; PFO: Upland Forest. Bold values indicate statistical significance at p value < 0.05. Distance index: Euclidean.

**Table S4.** Results of envfit analysis – correlation of the community structure with environmental variables for floodplain and forest areas.

| | Archaea | | Bacteria | |
|---|---|---|---|---|
| | $R^2$ | p value | $R^2$ | p value |
| *Chemical properties* | | | | |
| pH | 0.5683 | **0.001** | 0.6001 | **0.001** |
| OM | 0.7893 | **0.001** | 0.8587 | **0.001** |
| N | 0.7341 | **0.001** | 0.8673 | **0.001** |
| P | 0.0426 | 0.645 | 0.6797 | **0.001** |
| S | 0.0043 | 0.956 | 0.0720 | 0.471 |
| K | 0.0849 | 0.395 | 0.6660 | **0.001** |
| Ca | 0.1992 | 0.108 | 0.2627 | **0.039** |
| Mg | 0.1861 | 0.129 | 0.6308 | **0.001** |
| Al | 0.0195 | 0.812 | 0.4741 | **0.003** |
| B | 0.6436 | **0.001** | 0.6736 | **0.001** |
| Cu | 0.1919 | 0.111 | 0.7761 | **0.001** |
| Fe | 0.2272 | 0.069 | 0.0508 | 0.567 |
| Mn | 0.2656 | **0.034** | 0.7295 | **0.001** |
| Zn | 0.1754 | 0.119 | 0.5750 | **0.001** |
| *Factors* | | | | |
| Area | 0.8204 | **0.001** | 0.8817 | **0.001** |
| Season | 0.0031 | 0.913 | 0.0419 | 0.377 |

pH: hydrogen potential; OM: organic matter; N: nitrogen, P: phosphorus; S: sulphur; K: potassium; Ca: calcium; Mg: magnesium; Al: aluminium; B: boron; Cu: cupper; Fe: iron; Mn manganese; Zn: zinc. Bold values indicate statistical significance at p value < 0.05.

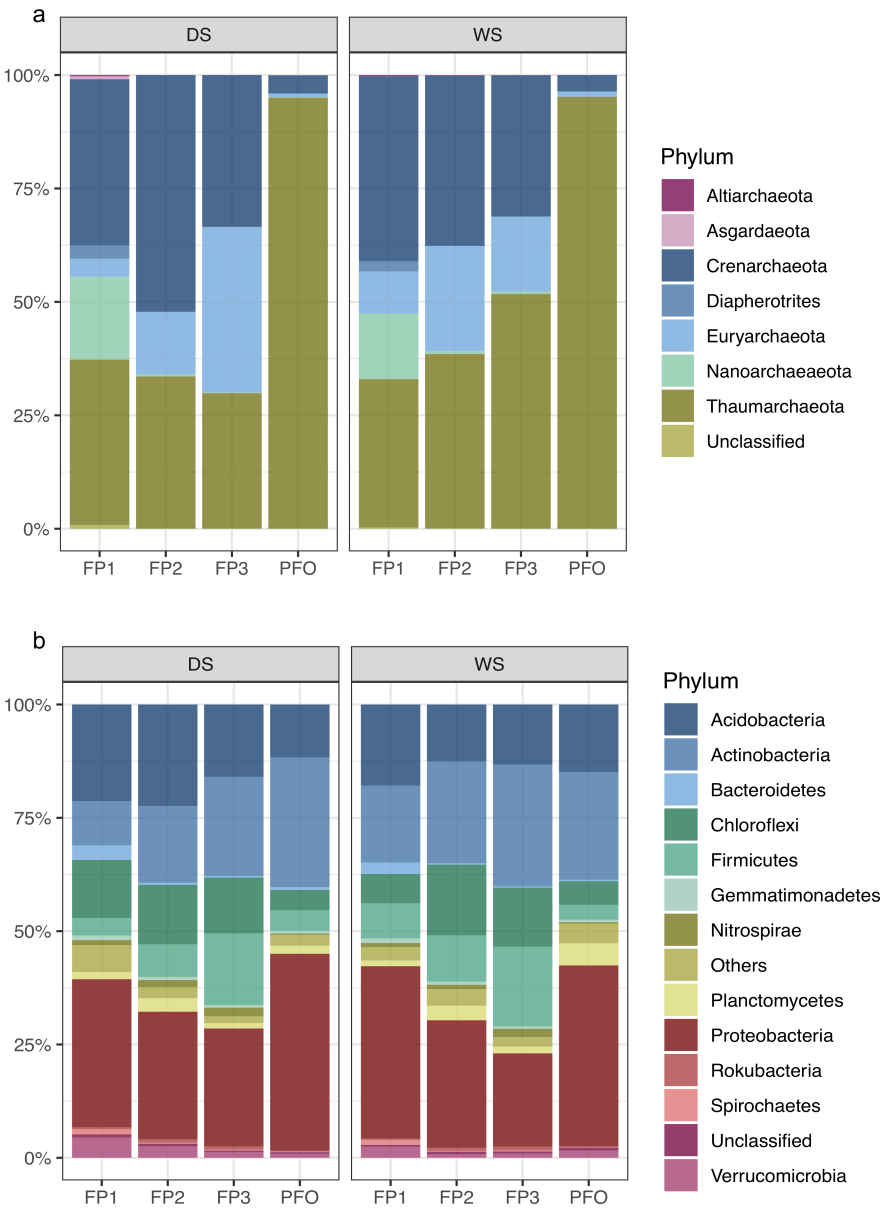


**Figure S1.** Relative abundance of archaeal (a) and bacterial (b) communities in the floodplain sediments (FP1, FP2 and FP3) and upland forest soils (PFO) during wet (WS) and dry (DS) seasons. Abundance values presented as an average of three replicates.

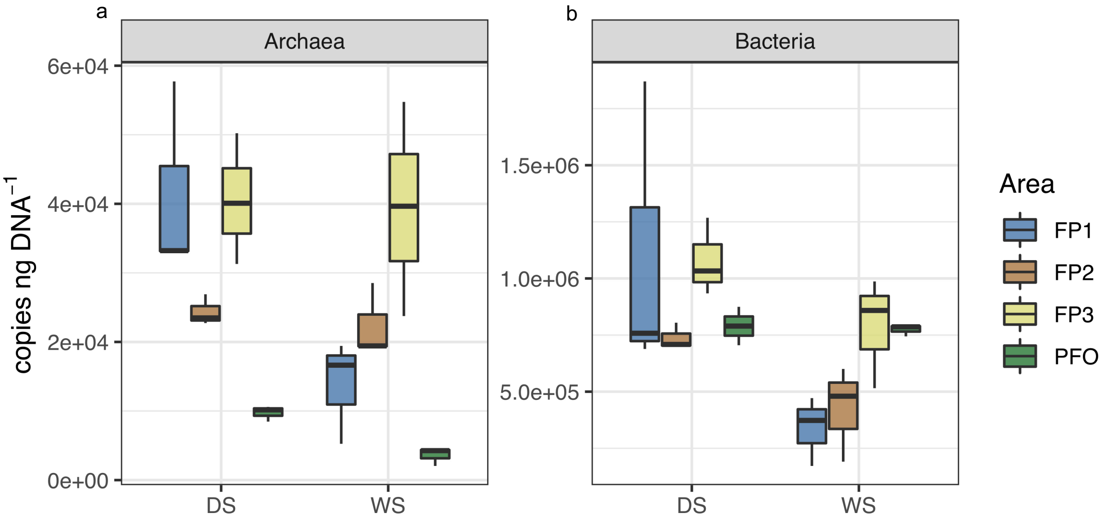


**Figure S2.** Number of copies per ng of DNA (copies ng $DNA^{-1}$) of archaeal (a) bacterial (b) 16S rRNA genes in the floodplain sediments (FP1, FP2 and FP3) and upland forest soils (PFO) during wet (WS) and dry (DS) season.
